# The effect of lysergic acid diethylamide (LSD) on whole-brain functional and effective connectivity

**DOI:** 10.1101/2022.11.01.514687

**Authors:** Peter Bedford, Daniel J. Hauke, Zheng Wang, Volker Roth, Monika Nagy-Huber, Friederike Holze, Laura Ley, Patrick Vizeli, Matthias E. Liechti, Stefan Borgwardt, Felix Müller, Andreea O. Diaconescu

**Author notes:** These authors contributed equally (Peter Bedford, Daniel J. Hauke, Felix Müller, Andreea O. Diaconescu). Corresponding author: Daniel J. Hauke, 250 College St., 12th Floor, Toronto, ON, M5T 1R8Canada.

## Abstract

Psychedelics have emerged as promising candidate treatments for various psychiatric conditions, and given their clinical potential, there is a need to identify biomarkers that underlie their effects. Here, we investigate the neural mechanisms of lysergic acid diethylamide (LSD) using regression dynamic causal modelling (rDCM), a novel technique that assesses whole-brain effective connectivity (EC) during resting-state functional magnetic resonance imaging (fMRI). We modelled data from two randomized, placebo-controlled, double-blind, cross-over trials, in which 45 participants were administered 100*μ*g LSD and placebo in two resting-state fMRI sessions. We compared EC against whole-brain functional connectivity (FC) using classical statistics and machine learning methods. Multivariate analyses of EC parameters revealed widespread increases in interregional connectivity and reduced self-inhibition under LSD compared to placebo, with the notable exception of primarily decreased interregional connectivity and increased self-inhibition in occipital brain regions. This finding suggests that LSD perturbs the Excitation/Inhibition balance of the brain. Moreover, random forests classified LSD vs. placebo states based on FC and EC with comparably high accuracy (FC: 85.56%, EC: 91.11%) suggesting that both EC and FC are promising candidates for clinically-relevant biomarkers of LSD effects.

## 1 Introduction

Psychedelics like psilocybin and lysergic acid diethylamide (LSD) have emerged as promising new treatment candidates for a variety of psychiatric conditions including alcohol and tobacco dependence (Rucker et al., 2018; Krebs and Johansen, 2012; Bogenschutz et al., 2022), as well as major depression and anxiety disorders (Griffiths et al., 2016; Ross et al., 2016; Gasser et al., 2015; Holze et al., 2022). Since psilocybin led to lasting positive changes in mood in healthy volunteers (Griffiths et al., 2006), it was recently investigated as a treatment for patients with depression (Carhart-Harris et al., 2016a, 2018; Davis et al., 2021). Carhart-Harris and colleagues presented first evidence that two doses of psilocybin may be as efficacious as a first-line antidepressant treatment (i.e., escitalopram), while being associated with far fewer side effects (Carhart-Harris et al., 2021). Since both LSD and psilocybin primarily act on 5-HT_2A_ receptors, an increasing number of clinical trials using psychedelics have been registered to evaluate their efficacy for treatment of depressive disorders and other psychiatric conditions (Nutt and Carhart-Harris, 2021). Given this clinical potential and growing interest in precision psychiatry, we sought to examine the neural mechanisms that underpin whole-brain effects of LSD by employing computational modelling and machine learning to evaluate the potential for individual-level predictions.

Functional connectivity (FC) (Tagliazucchi et al., 2016; Roseman et al., 2016; Kaelen et al., 2016; Carhart-Harris et al., 2016b; Müller et al., 2017; Preller et al., 2018; Barnett et al., 2020; Bershad et al., 2020; Luppi et al., 2021) and effective connectivity (EC) (Timmermann et al., 2018; Preller et al., 2019) have already shown promise in providing a framework for uncovering the neural mechanisms underlying LSD. Both connectivity measures are widely used, but are interpretatively distinct: FC is commonly assessed using Pearson correlation coefficients – a measure of the linear relationship – between the BOLD signal time series of two distinct brain regions. In a general linear model, the squared correlation coefficient represents the proportion of one signal’s variance, which can be explained by another signal, and vice versa. Though much of the research investigating FC changes under LSD suggests that FC is a promising candidate for a clinically-relevant biomarker (Tagliazucchi et al., 2016; Roseman et al., 2016; Kaelen et al., 2016; Carhart-Harris et al., 2016b; Müller et al., 2017; Preller et al., 2018; Barnett et al., 2020; Bershad et al., 2020; Luppi et al., 2021), FC can be limited in terms of its interpretability.

Firstly, FC is an undirected measure of connectivity, because computation of the correlation coefficient is commutative. Secondly, it ignores two important organizational principles of the cortex: (1) the asymmetry of connections (Felleman and Van Essen, 1991; Markov et al., 2014) and (2) the presence of gain regulation *within* a cortical region or ‘self-connections’. In contrast to FC, EC rests on a mechanistic model of how the data were generated (Friston, 1994) and estimates both, asymmetry and the gain within a cortical region. Here, we adopt a mechanistic perspective and examine LSD-driven changes in *directed* influences between nodes, as well as asymmetry and self-inhibition across the brain.

Here, we estimated whole-brain EC using regression dynamic causal modelling (rDCM; Frässle et al. (2017)), a recently developed variant of dynamic causal modelling (DCM; Friston et al. (2003)). rDCM allows estimation of whole-brain EC by applying several modifications and simplifications to the original DCM framework by reformulating a linear DCM in the time domain as a linear Bayesian regression in the frequency domain (Frässle et al., 2018). Furthermore, this model has been recently extended to allow modelling of task-free or resting-state magnetic resonance imaging (rs-MRI) data (Frässle et al., 2017, 2021b), enabling us to study – for the first time – how whole-brain effective connectivity is altered under LSD. Modelling asymmetry of directed influences and within-region gain can potentially provide additional useful information about the neural mechanisms underlying LSD effects.

To assess this, we also compared FC and EC in terms of their ability to distinguish LSD from placebo at the individual level. In future studies, these computationally-informed biomarkers could potentially be leveraged to predict the subjective effects of psychedelics, which have been associated with subsequent treatment response (Carhart-Harris et al., 2018; Roseman et al., 2018), to personalise clinical interventions for individual patients.

## 2 Results

### 2.1 The effect of LSD on functional connectivity

#### 2.1.1 Mass-univariate analysis

About 23% (1993/8646) unique correlation coefficients significantly differed across LSD and placebo conditions (*p* < 0.05). Among these connections, we observed widespread increases in FC under LSD (Figure 1A-C). Of the top 50 connections (ranked by t-statistic of the difference between conditions; Figure 2A), only two connections showed decreased FC under LSD, each connecting two regions in the occipital cortex: bilateral connections between occipital poles and the occipital fusiform gyri (ranked 6th and 8th respectively; Figure 2B). Six of the remaining top ten connected parietal and frontal regions (Figure 2A-B).

**Figure 1:**
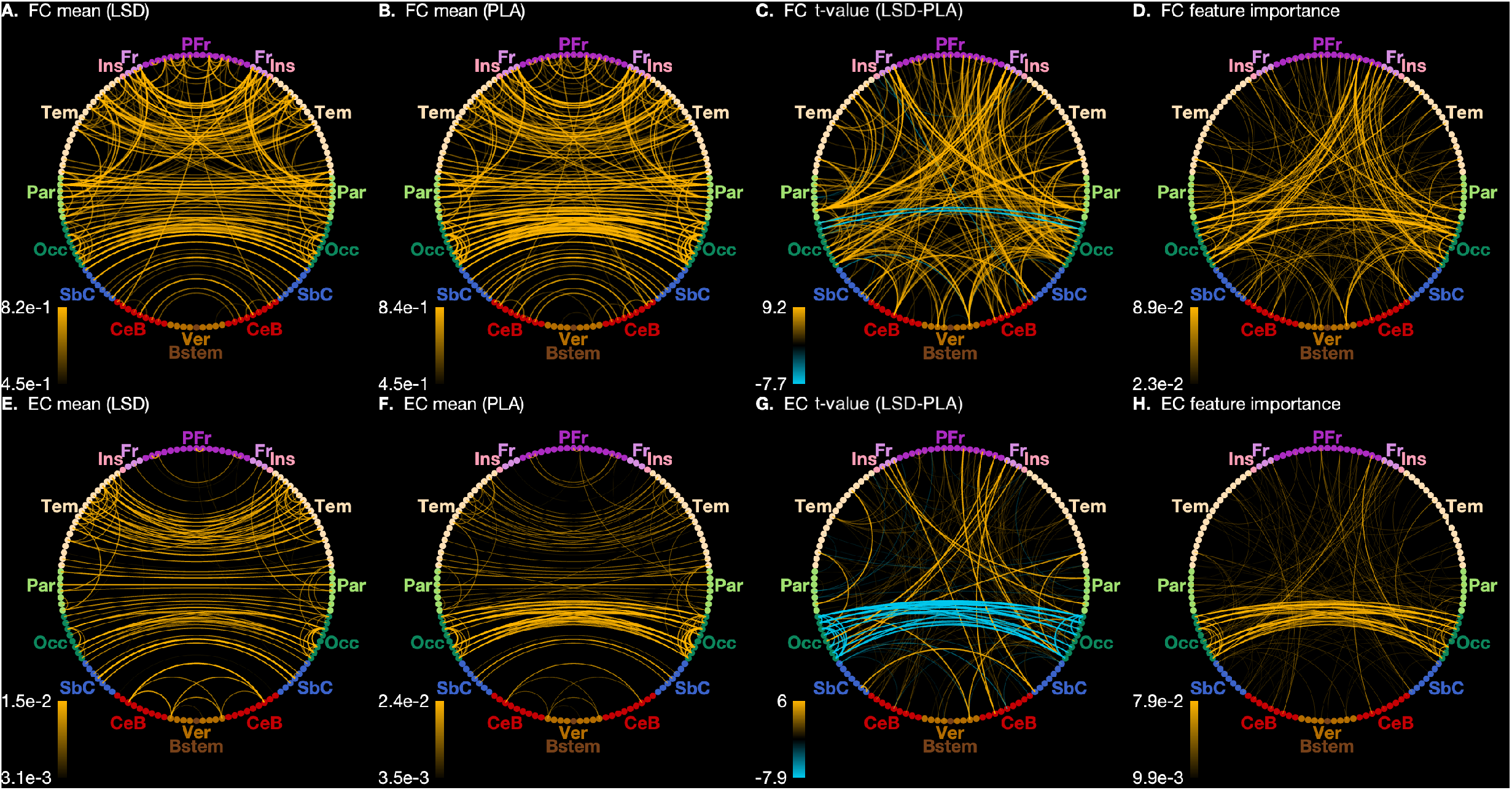
Connectogram views of differences in functional (FC) and effective connectivity (EC) between LSD and placebo conditions. **A** Across-participant average FC in the LSD condition. **B** Across-participant average FC in the placebo condition. **C** Across-participant í-statistic values of difference between FC in LSD and placebo conditions. **D** Feature importance estimates for the FC classification model. See section 4.4.3 for a detailed definition of feature importance. **E** Across-participant average EC in the LSD condition. **F** Across-participant average EC in the placebo condition. **G** Across-participant í-statistic values of difference between EC in LSD and placebo conditions. **H** Feature importance estimates for the EC classification model. Differences in magnitudes of connectivity are indicated in each connectogram by both line width and opacity. In **C** (FC) and **G** (EC), orange and blue lines indicate increases and decreases, respectively, in connectivity in the LSD condition. Note that for **E,F,G**, both directional EC values between each pair of regions have been averaged for display. To maintain visibility, only the top 250 connections have been displayed. PFr: Prefrontal cortex. Fr: Frontal cortex. Ins: Insular cortex. Tem: Temporal cortex. Par: Parietal cortex. Occ: Occipital cortex. SbC: Subcortical regions. CeB: Cerebellum. Ver: Vermis. Bstem: Brainstem.

**Figure 2:**
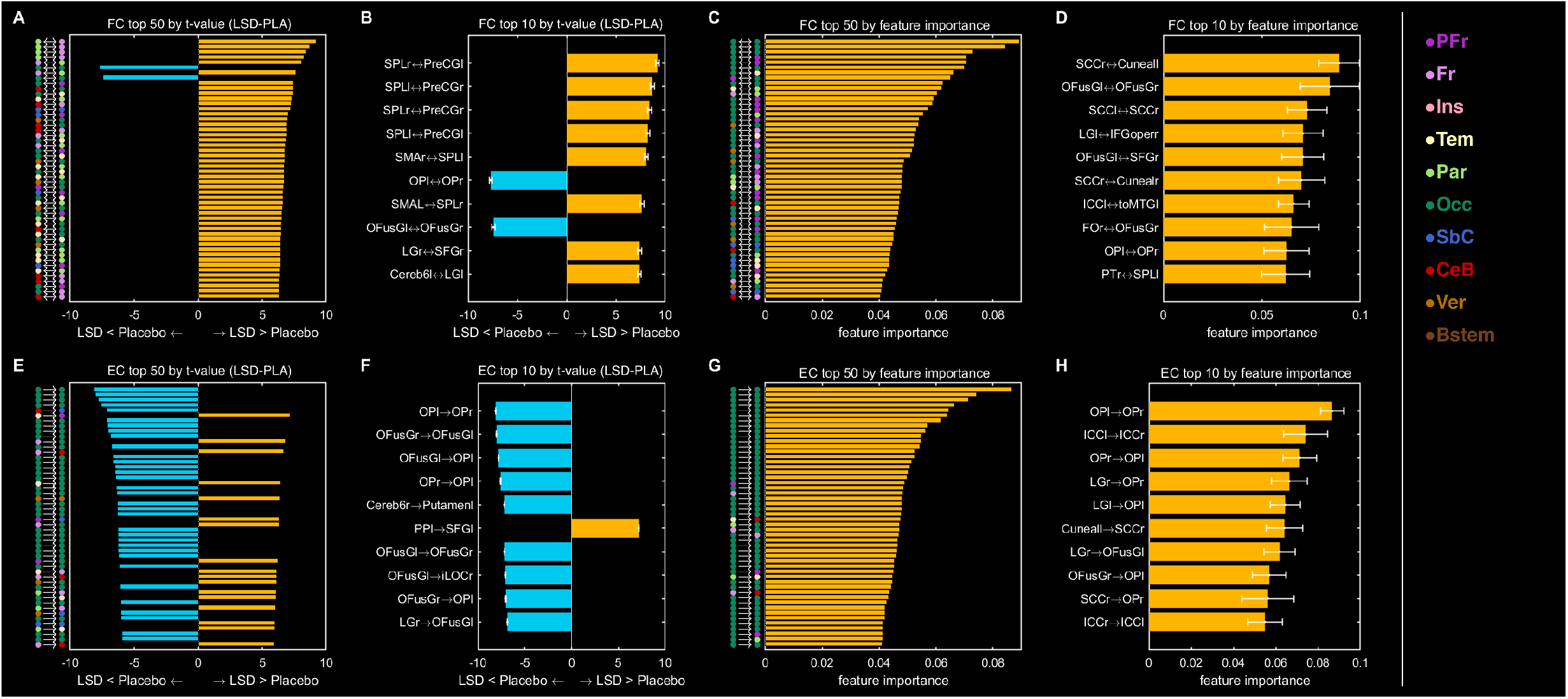
Ranked differences in functional (FC) and effective connectivity (EC) between LSD and placebo conditions. **A** Top 50 connections by across-participant t-statistic of the difference in FC between the LSD and placebo conditions. **B** Top 10 connections by across-participant t-statistic of the difference in FC between the LSD and placebo conditions. **C** Top 50 connections by feature importance estimate for the FC classification model. See section 4.4.3 for a detailed definition of feature importance. **D** Top 10 connections by feature importance estimate for the FC classification model. **E** Top 50 connections by across-participant t-statistic of the difference in EC between the LSD and placebo conditions. **F** Top 10 connections by across-participant t-statistic of the difference in EC between the LSD and placebo conditions. **G** Top 50 connections by feature importance estimate for the EC classification model. **H** Top 10 connections by feature importance estimate for the EC classification model. For **A,C,E,G**, coloured circles indicate the area into which each region of interest (ROI) is categorized. For **B,D,F,H**: orange and blue bars indicate increases and decreases, respectively, in connectivity in the LSD condition; abbreviations indicate the ROIs forming each connection. For **B,F**, errorbars represent the across-participant standard deviation of the differences in connectivity between conditions. For **D,H**, errorbars represent the across-fold standard deviation for the feature importance estimates. PFr: Prefrontal cortex. Fr: Frontal cortex. Ins: Insular cortex. Tem: Temporal cortex. Par: Parietal cortex. Occ: Occipital cortex. SbC: Subcortical regions. CeB: Cerebellum. Ver: Vermis. Bstem: Brainstem.

#### 2.1.2 Partial least squares correlation analysis

PLSC analysis of FC showed a significant condition effect (LSD condition score: 4.76 [4.11, 5.39], p < 0.001). LV loadings indicated that FC was stronger under LSD compared to placebo across a large number of regions. The most reliable effects were observed for the following connections: right thalamus and left hippocampus, bilateral anterior cingulate cortex (ACC) and bilateral intracalcarine cortex, left and right dorsolateral PFC, right superior frontal gyrus and right lingual gyrus, right frontal pole and right thalamus, and right middle frontal gyrus and right lingual gyrus (bootstrap ratios (BSR) > 6.4). Conversely, increased FC under placebo was found between several occipital regions, including left and right fusiform gyrus, supracalcarine cortex, occipital pole, and between bilateral putamen and cerebellum (BSR ≤ −5.0;Figure 4).

**Figure 3:**
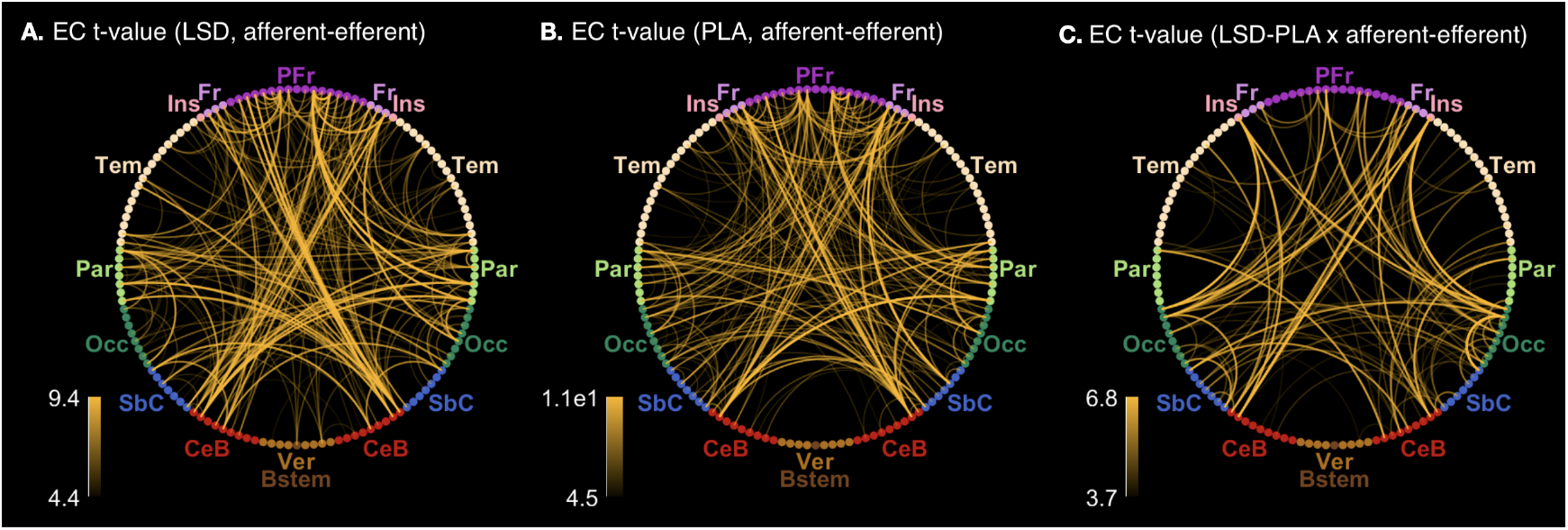
Connectogram views of asymmetries in effective connectivity (EC). **A** Across-participant t-statistic of the difference in EC between the two directions of influence between each pair of regions, for the LSD condition. **B** Across-participant t-statistic of the difference in EC between the two directions of influence between each pair of regions, for the placebo condition. **C** Across-participant t-statistic of the difference in EC between the two directions of influence between each pair of regions, and between the LSD and placebo conditions. Differences in magnitudes of connectivity and connectivity changes are indicated in each connectogram by both line width and opacity. To maintain visibility, only the top 250 connections have been displayed. PFr: Prefrontal cortex. Fr: Frontal cortex. Ins: Insular cortex. Tem: Temporal cortex. Par: Parietal cortex. Occ: Occipital cortex. SbC: Subcortical regions. CeB: Cerebellum. Ver: Vermis. Bstem: Brainstem.

**Figure 4:**
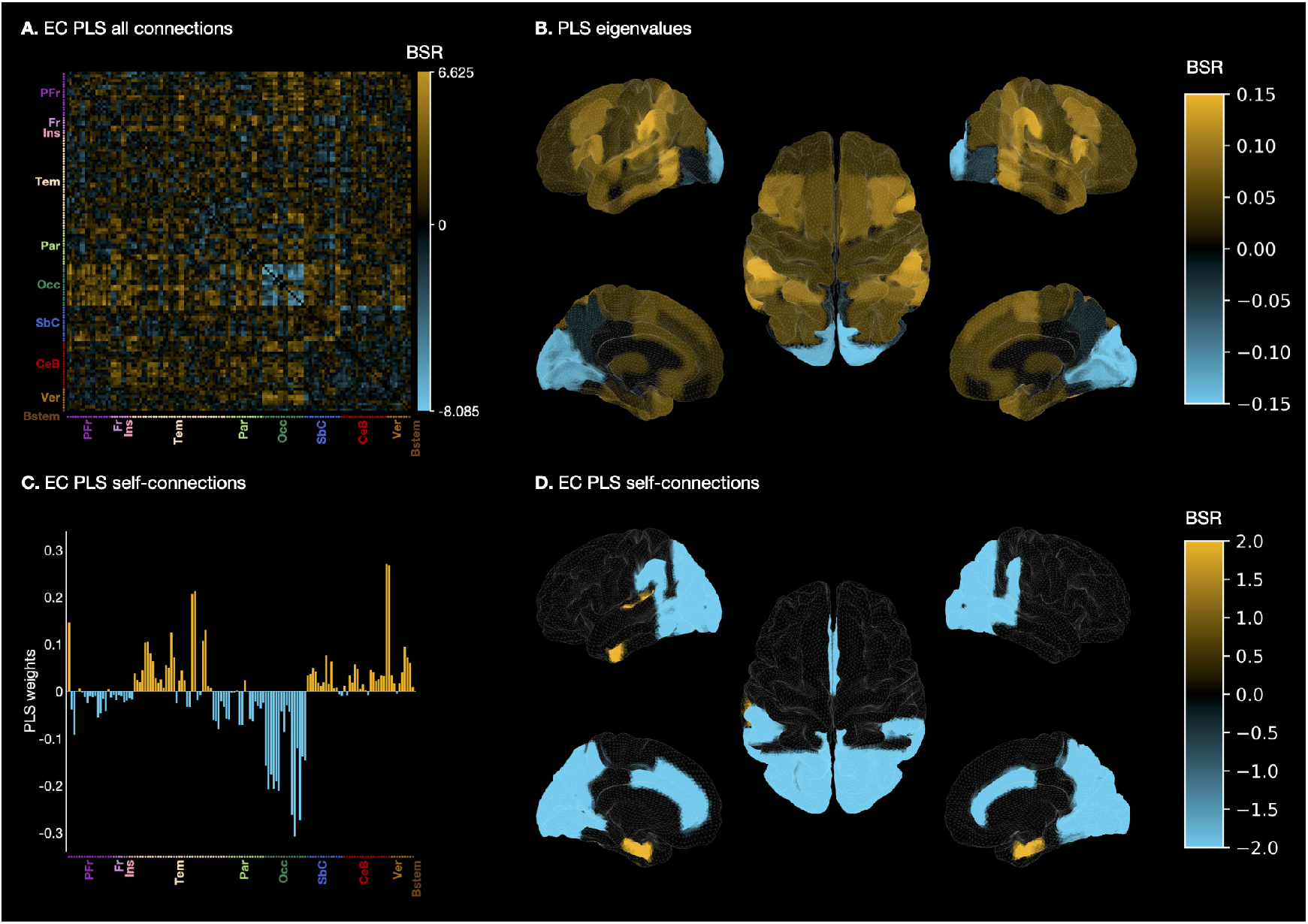
Graphical and anatomical visualizations of partial least squares (PLS) correlation analysis results. **A** Whole-brain EC BSRs (bootstrap ratios) reflecting condition differences **B** Leading Eigenvector reflecting condition differences in whole-brain EC across brain regions **C** Brain region saliences reflecting condition differences across self-connections **D** Brain region BSRs reflecting condition differences across self-connections For A-D: orange and blue areas indicate increases and decreases, respectively, in connectivity in the LSD compared to placebo conditions. PFr: Prefrontal cortex. Fr: Frontal cortex. Ins: Insular cortex. Tem: Temporal cortex. Par: Parietal cortex. Occ: Occipital cortex. SbC: Subcortical regions. CeB: Cerebellum. Ver: Vermis. Bstem: Brainstem. BSR: Bootstrap ratio.

#### 2.1.3 Machine learning analysis

The random forest trained on FC as features (FC model) discriminated between LSD and placebo with a BAC of 86% (*p* < 0.001; Table 1). Feature importance measures suggested that connections involving occipital and prefrontal regions were most relevant for this classification performance. Of the top 50 connections (ranked by feature importance), only twelve did not involve occipital regions. Of those twelve, four connected frontal and parietal regions, and the remaining temporal and parietal regions (Figure 2C). The connections remaining in the top 10 involved connections between occipital brain regions as well as connections from occipital regions to prefrontal and temporal regions (Figure 2D).

**Table 1:** Classification Performances. Performance measures for models trained on either functional connectivity (FC) or effective connectivity (EC). *N_tree_:* Number of trees used in the random forest models. BAC: Balanced accuracy. AUC: Area-under-the-curve. SE: Sensitivity. SP: Specificity. PPV: positive predictive value. NPV: negative predictive value. ^*a*^ p-values for balanced accuracy were obtained through permutation test with 1000 label permutations.

| | $N_{tree}$ | BAC | AUC | SE | SP | PPV | NPV | $p\text{-value}^a$ |
| --- | --- | --- | --- | --- | --- | --- | --- | --- |
| FC | 8646 | 0.86 | 0.95 | 0.87 | 0.84 | 0.85 | 0.87 | $<0.001$ |
| EC | 17424 | 0.91 | 0.96 | 0.91 | 0.91 | 0.92 | 0.92 | $<0.001$ |

### 2.2 The effect of LSD on effective connectivity

#### 2.2.1 Mass-univariate analysis

About 13% (2184/17424) effective connections strength coefficients significantly differed across conditions (*p* < 0.05). As with LSD-induced changes in FC, we observed widespread increases in EC under LSD (Figure 1E-G). However, surprisingly, among the most prominent differences were *decreases* in EC between occipital brain regions under LSD, including 33 out of the top 50 (Figure 2E) and nine out of the top 10 connections (Figure 2F) ranked by t-statistic of the difference between conditions. Only 13 out of the top 50 connections did not involve an occipital region. Of the top 10 connections, only 2 were not between occipital regions. These included right cerebellum to left putamen, which showed reduced EC, whereas left planum temporale-to-left superior frontal gyrus connections showed increased EC under LSD compared to placebo (Figure 2F).

#### 2.2.2 Partial least squares correlation analysis

PLSC analysis of EC showed a statistically significant condition effect (LSD condition score: 0.043 [0.047, 0.037], *p* < 0.001). Similar to the PLSC results based on FC data, we found that EC was primarily increased under LSD. Reliable loadings were observed for connections between (i) occipital and prefrontal regions, including bilateral occipital pole to bilateral ACC, bilateral fusiform gyrus to bilateral ACC, right lingual gyrus to right dorsolateral PFC and right dorsolateral PFC to right lingual gyrus, (ii) right thalamus to right frontal pole and right frontal pole to right thalamus, and (iii) occipital and temporal regions, including the right lateral occipital cortex to right superior temporal gyrus and the left middle temporal gyrus to left intracalcarine cortex (BSR > 5.5). The reverse pattern showing decreased EC under LSD was observed exclusively between the left and the right hemispheres of several occipital regions, including the occipital pole, lingual gyrus, supracalacarine cortex, and fusiform gyrus (BSR < −5.5; Figure 4).

#### 2.2.3 Machine learning analysis

The random forest trained on EC as features (EC model) performed with a BAC of 91.1% (*p* < 0.001; Table 1). As with the mass-univariate and PLSC analysis, connections involving occipital regions were the most predominant features driving the classification performance (Figure 2G-H). Of the top 50 connections (ranked by feature importance), the majority (39/50) represented connections between pairs of occipital regions. The highest-ranked connection invovling a non-occipital region included right dorsolateral PFC to right lingual gyrus and left dorsolateral PFC to left cerebellum. This finding suggests that these connections were not only significantly different under LSD (see Mass-univariate analysis section), but also contributed substantially to highly-accurate individual-level predictions.

### 2.3 Comparing functional vs effective connectivity changes under LSD

### 2.4 The effect of LSD on inhibitory self-connections

As pointed out in the Introduction, an advantage of using EC over FC is that rDCM allows estimation of inhibitory self-connections, or the coefficients along the diagonal of the **A** matrix. These all-negative values can be interpreted as ‘decay-rate’ coefficients (in view of the DCM state equation) since they capture the tendency of different regions to return to baseline, rather than increasing activity indefinitely. Also note that these values reflect local as opposed to global (inter-regional) dynamics.

#### 2.4.1 Mass-univariate analysis

About 30% (39/132) inhibitory self-connections significantly differed across conditions (p < 0.05). Occipital regions once again displayed the greatest effects showing more negative or decreased local connectivity under LSD (Figure 5). One exception to this pattern was an increase for the Vermis 8 self-connection, which ranked second in terms of absolute t-value. The top-ranked self-connection was the left occipital pole, which was involved in the first-, third-, fourth-, and ninth-ranked non-self-connections by absolute t-value.

**Figure 5:**
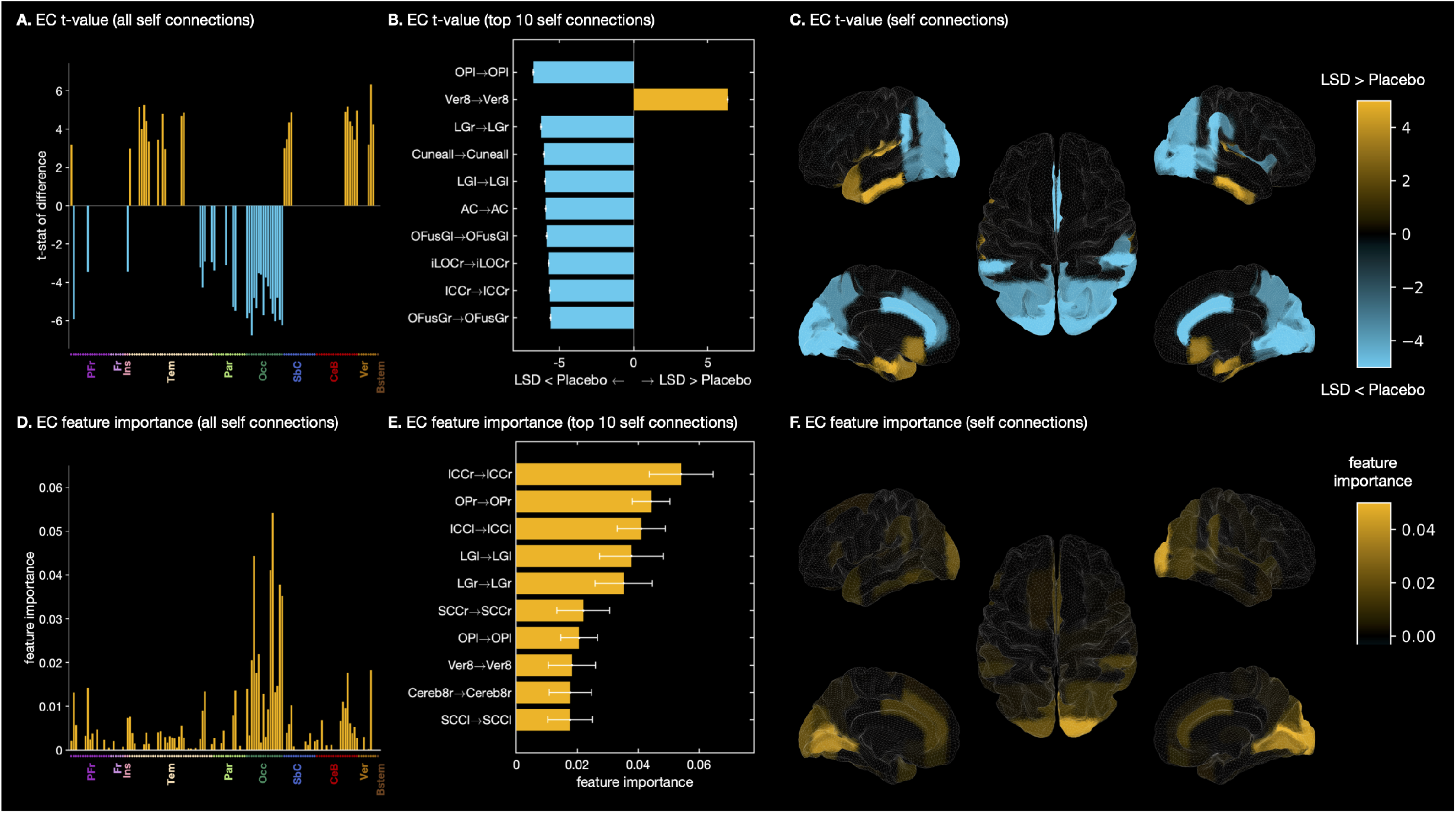
Effect of LSD on inhibitory self-connections. **A** t-statistic of the difference between LSD and placebo conditions in self-connections. **B** Top 10 self-connections ranked by t-statistic of the difference between LSD and placebo conditions. **C** Anatomical colourmap of t-statistic of the difference between LSD and placebo conditions in self-connections. **D** Estimates of feature importance of self-connections in EC classification model. **E** Top 10 self-connections by feature importance in the EC classification model. **F** Anatomical colourmap displaying feature importance of self-connections in the EC classification model. For **A,B,C**: orange and blue areas indicate increases and decreases unde LSD, respectively. For **B**, errorbars represent the across-participant standard deviation of the differences in connectivity between conditions. In **B** and **E**, abbreviations indicate the ROIs forming each connection. For **E**, errorbars represent the across-fold standard deviation of the feature importance estimates. PFr: Prefrontal cortex. Fr: Frontal cortex. Ins: Insular cortex. Tem: Temporal cortex. Par: Parietal cortex. Occ: Occipital cortex. SbC: Subcortical regions. CeB: Cerebellum. Ver: Vermis. Bstem: Brainstem.

#### 2.4.2 Partial least squares correlation analysis

LSC analysis performed on self-connections showed a significant condition effect (LSD condition score: −0.043 [−0.047, −0.029], p < 0.001). Self-connections were more inhibitory (i.e., became more negative) under LSD primarily in occipital, parietal, and prefrontal regions (Figure 4C4). The largest increases in self-inhibition under LSD occurred in the bilateral lingual gyri, fusiform gyri, occipital pole, and supracalcarine cortex (Figure 4D4). Conversely, disinhibition under LSD was observed in temporal, subcortical and brainstem regions (Figure 4C4). The largest disinhibition under LSD was noted in the bilateral hippocampi and parahippocampal gyri, globus pallidus, and the cerebellum (Figure 4D4).

Note that all self connections are negative. Disinhibition in any individual brain area suggests that this area becomes less stable under LSD from a dynamic system viewpoint, since the system moves closer to a critical point as self-connections approach zero.

#### 2.4.3 Machine learning analysis

Ranking self-connections by feature importance, we found again that many of the most important connections were occipital. Among the top 10 self-connections ranked by feature importance were two non-occipital connections, namely Vermis 8 and right Cerebellum 8 (Figure 5).

### 2.5 Asymmetry in directed connectivity

Recall that while FC provides a measure of the correlation between the activities of each pair of brain regions, EC provides an estimate of directed effects between pairs of regions, which may be asymmetrical. We tested for the presence of asymmetry (by mass-univariate comparison) in each condition (LSD and placebo), and tested whether asymmetry was affected by LSD (drug-by-asymmetry interaction). The number of forwardbackward pairs of endogenous connectivity coefficients that were significantly different (*p* < 0.05) was 868 out of 8648 (10%) under LSD, 1202 (14%) in the placebo condition, and 120 (1.4%) for the drug-by-asymmetry interaction.

### 2.6 Comparing functional and effective connectivity classifiers

Lastly, we compared the performances of classifiers trained on either FC or EC to test whether modelling additional physiological details (i.e., self-inhibition and asymmetry) translated into a better classification performance of drug condition. While BACs differed numerically between the FC and EC classification models (86% vs 91% BAC), a McNemar’s test comparing the models’ performances suggested that these differences were not significantly different (p > 0.074), indicating that both FC and EC may be promising biomarkers for individual-level predictions.

## 3 Discussion

The goal of this study was to investigate the effects of LSD on whole-brain EC and gauge its potential as a biomarker for clinically-relevant predictions. To this end, we studied LSD-induced effects on whole-brain EC through mass-univariate, multivariate, and machine learning analyses and compared EC to FC. Mass-univariate analysis revealed global increases in both FC and EC under LSD, with the notable exception of connections involving occipital regions, which decreased. Multivariate PLSC analyses confirmed that EC between regions in occipital cortices decreased under LSD, while EC between parietal, temporal and inferior frontal regions was increased. When comparing FC to EC, we found notable changes in self-inhibition in around 30% of the brain regions under LSD, indicating that LSD may perturb the excitation/inhibition (E/I) balance of the brain. Moreover, our results suggest that – while EC was asymmetric (10-14% of connections) – asymmetry was largely unaffected by LSD (1.4% of connections). Lastly, our machine learning analysis showed that both FC and EC constitute promising biomarkers for future individual-level predictions of clinically-relevant targets, since classifiers trained on either FC or EC could discriminate between LSD and placebo with high accuracy of 86% and 91%, respectively.

### 3.1 Face validity of whole-brain effective connectivity

To assess the face validity of EC measures, we compared thalamic connections and found that the effects (sign and significance) were consistent across FC and EC, although a recent study argued that this does not necessarily need to be the case (Barnett et al., 2020). Indeed, changes in thalamic connectivity across conditions were also in perfect agreement with those of a previous FC analysis based on a subset participants (Müller et al., 2017). Together with the highly accurate classification performance, these findings reassured us that EC estimates were plausible.

Comparing all connections, we found that FC and EC measures were broadly in agreement with one another as both indicated generally increased connectivity under LSD, with some notable connectivity decreases in bilateral occipital areas, though this effect was more pronounced in the EC measure. Feature importance also suggested that the EC classification relied more heavily on the bilateral occipital connections than the FC classification, wherein connections involving more varied areas were represented among the highlighted ‘important’ features (Figure 2).

### 3.2 Possible interpretation of occipital effects

A striking aspect of our results is that LSD appears to have a different effect on EC in occipital areas compared to most other brain regions. Changes in these connections may underlie subjective LSD effects that involve visual hallucinations in particular, as Kaelan et al. (2016) (Kaelen et al., 2016) found that increased parahippocampus to visual cortex EC correlated with enhancements in eyes-closed mental-imagery during music-listening under LSD.

Moreover, Roseman et al. (2016) found that activity within visual cortex becomes more dependent on its intrinsic retinotopic organization under LSD, and posit that the early visual cortex appears to behave as if the subjects were seeing external visual inputs in the eyes-closed state. It is well-known that occipital alpha oscillations are strongest in the eyes-closed state or in the absence of visual stimulation (Nierhaus et al., 2009); LSD-induced widespread decreases in occipital connectivity in the eyes-closed state may be another reflection of eyes-open-like-behaviour of the visual system. Indeed, ratings of ‘simple hallucinations’ under LSD were found to correlate with increased V1 resting state FC and decreased alpha power (Carhart-Harris et al., 2016b).

### 3.3 Comparison with other studies investigating directed connectivity

Unlike another study that investigated directed connectivity using Granger causality based on magnetoencephalography (MEG) recordings (Barnett et al., 2020), we found predominantly increased EC mirroring increased FC patterns across the brain with the exception of the aforementioned occipital connections. This difference may be explained by the different methods used (Granger causality measuring directed FC vs rDCM measuring EC) or the measurement modalities and resulting differences in temporal resolution (MEG vs fMRI).

Moreover, Preller et al. (2019) investigated EC in cortico-striato-thalamo-cortical feedback loops in a sparse network to test the thalamic gating hypothesis using spectral DCM. Their chosen network consisted of the thalamus, ventral striatum (VS), posterior cingulate cortex (pCC), and temporal cortex. In line with their results, we found that thalamus → pCC connectivity increased under LSD, while the finding that the VS → thalamus connection decreased was not reproduced in our study. Only the right accumbens → right thalamus connection was increased under LSD. However, the VS → thalamus decrease was independent of the Ketanserin condition, while the thalamus → pCC increase was modulated by 5-HT_2A_ receptor activation – perhaps pointing to differing mechanisms of action.

### 3.4 Implications for the thalamic gating hypothesis and brain entropy accounts

Our results are in line with the thalamic gating hypothesis, which postulates that psychedelics may temporally reduce thalamic gating leading to excessive information flow from thalamus to cortical regions (Vollenweider and Geyer, 2001). As previous analyses investigating FC (Müller et al., 2017; Tagliazucchi et al., 2016) and EC (Preller et al., 2019) changes under LSD, our results suggest that connectivity from thalamus to a widespread network of cortical regions is increased.

Brain entropy accounts of psychedelics propose that altered states of consciousness observed following administration of psychedelics result from increased entropy in the brain (Carhart-Harris et al., 2014; Carhart-Harris, 2018; Carhart-Harris and Friston, 2019). Even though testing these accounts was not the goal of this study, we note that overall our results are in agreement with this proposal. Specifically, we found changes in self-connections or within-region dynamics suggesting widespread disinhibition across most of the cortex. From a dynamic system perspective, this disinhibtion renders the system more unstable since it approaches a critical point as self-connection values approach zero. However, it is worth noting that we found the opposite pattern in occipital regions, where local inhibition increased under LSD rendering dynamics in these areas more stable. While our results are generally in line with the proposal that psychedelics like LSD increase brain entropy (on average), this whole-brain analysis revealed that a more fine-grained assessment of entropy across regions may be warranted in future studies to refine these accounts as specifically occipital regions may be impacted differently.

It is interesting to note that connections between occipital regions appear to dominate EC under placebo (Figure 1F), whereas the difference between visual and other regions is reduced under LSD (Figure 1E). This result aligns with a recent study (Girn et al., 2022), which showed that the principal gradient of cortical connectivity flattens under LSD. Moreover, Girn et al. (2022) (Girn et al., 2022) do not only suggest that LSD reduces processing steps between unimodal and transmodal cortices, but point out further that their findings are in line with the RElaxed Beliefs Under Psychedelics (REBUS) model (Carhart-Harris and Friston, 2019). Our results support this interpretation of a decrease in functional differentiation between sensory and abstract cognitive processing under LSD and therefore appear to align with reports of a blurred internal-external/subject-object distinction and increased connection between mentation and perception under LSD.

### 3.5 Does LSD perturb the excitation/inhibition (E/I) balance?

Recent studies have begun to investigate the impact of psychedelics on glutamate-mediated excitation of the cortex. At least two different pathways for glutamatergic effects have been proposed: (1) Agonism at 5-HT_2A_ receptors on pyramidal cells may lead to increased glutamate release (Nichols, 2016; Aghajanian and Marek, 1999) and (2) agonism at postsynaptic, inhibitory 5-HT_1A_ receptors may result in reduced excitation in certain regions, for example in hippocampus (Lewis et al., 2017; Mason et al., 2020). While we cannot directly speak to these pathways, widespread increases in self-connections or local disinhibition as identified in this study may relate to 5-HT_2A_-mediated glutamate release. Conversely, increased inhibition could be mediated via the 5-HT_1A_ pathway, although this receptor is not strongly expressed in occipital regions (Beliveau et al., 2017). Rather, 5-HT_1B_ expression is increased in these regions (Beliveau et al., 2017) and may be a candidate mechanism requiring further investigation.

Our results along with preclinical studies (Aghajanian and Marek, 1999; Scruggs et al., 2000, 2003) and recent magnetic resonance spectroscopy results (Mason et al., 2020) suggest that LSD may perturb the E/I balance of the brain. This is especially relevant because disturbances in the E/I balance have been discussed in the context of psychosis (Jardri and Deneve, 2013; Jardri et al., 2017) and more recently in the context of psychedelic-induced hallucinations and synaesthesia (Leptourgos et al., 2022). Comparing LSD-induced changes in E/I balance to other psychedelics and those associated with clinical psychosis will be an interesting avenue for future research.

### 3.6 Limitations

A few limitations of this study merit attention. Firstly, McCulloch and colleagues (2022) (McCulloch et al., 2022) recently argued that cross-over designs may not be well suited to study LSD due to potential long term effects of psychedelics. However, arguably, carryover effects would diminish, rather than increase, the difference between conditions in this study.

Secondly, LSD is known to affect heart rate, blood pressure and body temperature (Schmid et al., 2015; Dolder et al., 2018). Here, we controlled for physiological effects using baseline measures. Nonetheless, these effects are likely still impacting connectivity estimates and may partially account for the effects. Future studies should include concurrent physiological measurements during MRI acquisition to allow for more detailed physiological noise modelling (e.g., see (Kasper et al., 2017)).

Thirdly, due to the salient subjective effects of LSD, blinding is inherently difficult. Thus, knowledge about the condition could have impacted neural effects and future studies should include active control conditions. Although do note that one of the studies included an active control (Holze et al., 2020), rendering it less likely that participants correctly identified the condition in part of the data.

Finally, while classification performances in this study were promising, they should be taken as preliminary until replicated in an external sample.

### 3.7 Future directions

Our results suggest that local gain is changed under LSD implicating disturbances of the E/I balance as a neural mechanism underlying LSD effects. Because rDCM summarises region-specific E/I balance by a single parameter per region, we cannot determine whether excitation, inhibition, or both are impacted. Future studies should employ more detailed models that allow to pinpoint these changes and compare them empirically to other conditions in which the E/I balance is affected, for example psychosis (Jardri et al., 2017; Leptourgos et al., 2022).

Furthermore, our results suggest that visual regions may be impacted quite differently both in terms of between-region connectivity as well as in terms of local gain compared to the rest of the cortex warranting further investigation. The physiological basis of these changes and the relationship with subjective effects – visual hallucinations in particular - should be examined in future studies.

Finally, we found that both whole-brain FC and EC were equally capable of discriminating between LSD and placebo with high accuracy suggesting that both are promising candidates for more challenging prediction targets. More research is needed to assess whether machine learning models trained on either FC and EC estimates are able to predict subjective effects of LSD at an individual level both in healthy controls and in clinical trials to adapt psychedelic-assisted therapy to the needs of individual patients.

### 3.8 Conclusions

To the best of our knowledge, this is the first study to examine the impact of LSD on whole-brain EC. In particular, we found that compared to placebo, LSD impacted local gain and induced widespread increases in whole-brain functional and EC with the notable exception of connections involving occipital regions. Both connectivity measures performed exceptionally as features in predicting the experimental conditions in a machine learning-based analysis highlighting the potential of using connectivity measures to predict subjective effects of LSD or LSD treatment outcomes in the future.

## 4 Materials and Methods

### 4.1 Participants

Data was aggregated across two randomized, placebo-controlled, double-blind, cross-over trials. We will briefly reiterate the most important aspects pertaining to our analysis below. Please, refer to the original publications for further details (trial A: (Dolder et al., 2016; Müller et al., 2018); trial B: (Holze et al., 2020)). Both studies were approved by the Ethics Committee for Northwest/Central Switzerland (EKNZ) and by the Federal Office of Public Health. All participants provided written consent prior to participating and received monetary compensation.

#### 4.1.1 Sample size and recruitment

Participants were recruited from the University of Basel through advertisement and word of mouth. In trial A, 20 healthy participants (10 male, 10 female; age 32 ± 11 [mean ± SD]; range 25 – 60 years; body weight, 68.8 ± 7.7*kg*) were selected out of initially 24 participants (*N* = 4 participants were excluded from analysis due to head motion > 2*mm* translation or > 2° rotation; see Section 4.4.1). Only two participants had used a hallucinogenic drug before, and both on only a single occasion. In trial B, 28 healthy participants (14 men, 14 women; age 28 ± 4; range 25 – 45 years; body weight, 71.5 ± 12.0kg) were recruited. Five participants had previously used a hallucinogen, including LSD (three participants, 1-4 times). Eight participants had never used any illicit drugs with the exception of cannabis. We analyzed data from a total of 45 participants collected from Trials A and B.

#### 4.1.2 Inclusion and Exclusion criteria

Participants between the ages of 25-26 years (trial A) and 25-50 years (tiral B) were included in the study. Exclusion criteria for both studies were: Age < 25, pregnancy, personal or first-degree relative history of major psychiatric disorders (assessed by the Semi-structured Clinical Interview for Diagnostic and Statistical Manual of Mental Disorders, 4th edition, Axis I disorders by a trained psychiatrist), tobacco smoking > 10 cigarettes/day, use of medications that may interfere with study medications (e.g. antidepressants, antipsychotics, sedatives), lifetime prevalence of illicit drug use > 10 times (excluding Δ^9^-tetrahydrocannabinol), illicit drug use within the last two months, and illicit drug use during the study (determined by urine drug tests).

Study-specific exclusion criteria included: Age > 65, history of drug dependence and nursing, cardiac or neurological disorders, hypertension (> 140/90 mmHg) or hypotension (SBP< 85 mmHg) in trial A and age > 50, chronic or acute physical illness (abnormal physical exam, electrocardiogram, or hematological and chemical blood analyses) in trial B. Both studies included general health assessments.

### 4.2 Experimental procedure

Participants were administered 100*μ*g LSD orally in capsules (trial A) or vials (trial (B) and identical mannitol and ethanol-filled placebo capsules/vials in a cross-over design across two separate experimental sessions with a time between sessions of at least 7 days (24.5 ± 19.6 days). Each session included assessment of brain activity during rest using fMRI. Participants were instructed to close their eyes and remain awake during the scan.

### 4.3 Data acquisition

All images were acquired using a 3 Tesla MRI system (Magnetom Prisma, Siemens Healthcare, Erlangen, Germany) with a 20-channel phased array radio frequency head coil. For anatomical images a T1-weighted MPRAGE sequence (field-of-view: 256 × 256*mm*^2^; resolution: 1 × 1 × 1*mm*^3^; TR: 2000*ms*; TE: 3.37*ms*; flip angle: 8°; bandwidth: 200*Hz*/pixel) was used. Resting-state fMRI was acquired using interleaved T2*-weighted EPI using the following parameters: 35 axial slices, slice thickness: 3.5mm, inter-slice gap: 0.5mm, field-of-view: 224 × 224*cm*^2^, resolution: 3.5 × 3.5 × 3.5*mm*^3^, TR: 1800*ms*, TE: 28*ms*, flip angle: 82°, and bandwidth: 2442*Hz*/pixel. Three hundred volumes were acquired (total acquisition time: 9*min*). This resting state scan was followed by other sequences not relevant for this publication. Total acquisition time for all sequences was approximately 50min. No significant differences in head motion were present between conditions (see (Müller et al., 2018) for more details).

### 4.4 Data analysis

All analyses were implemented in Matlab (versions used for preprocessing: 2019b and connectivity analyses: 2018a; https://mathworks.com/). We will report relevant functions and toolboxes for sub-analyses below.

#### 4.4.1 Preprocessing

Structural and functional neuroimaging data were preprocessed using the CONN toolbox (version: 19c; (Whitfield-Gabrieli and Nieto-Castanon, 2012); https://web.conn-toolbox.org/) based on SPM12 ((Friston et al., 2007); https://www.fil.ion.ucl.ac.uk/spm/). The first five volumes of the BOLD time series were discarded to ensure signal equilibrium. Preprocessing included slice-timing correction, realignment to the mean image, and co-registration to the participant’s own structural scan. The structural image underwent a unified segmentation procedure combining segmentation, bias correction, and spatial normalization (Ashburner and Friston, 2005). The same normalization parameters were then applied to the functional images. Functional images were smoothed with an isotropic Gaussian kernel (6*mm* FWHM).

Quality control assessments comprised three stages: First, all scans were assessed considering the percentage of scrubbed volumes. Participants were excluded if < 5min of the scan remained after scrubbing (corresponding to < 55% of the initial volumes). This was due to evidence indicating that resting-state scans < 5*min* are not reliable (Birn et al., 2013). No participant was excluded based on this criterion. Second, maximal head motion after the scrubbing procedure were assessed using frame-wise displacement (FD) (Power et al., 2012). A sphere radius of 50*mm* was chosen for the calculation, and participants were excluded if maximum FD was > 1.75*mm* (half-voxel size). Five participants were excluded based on this criterion, resulting in a final sample of 45 participants. The mean percentage of the scrubbed volumes for this sample was 5.5 ± 3.7% for the drug condition and 3.8 ± 2.6% for the placebo condition. Average maximum FD was 0.74 ± 0.39*mm* for the drug condition and 0.49 ±0.24*mm* for the placebo condition.

Further noise correction of the functional images included linear regression of the six motion parameters and the white matter and cerebrospinal fluid signals, using individual tissue masks obtained from the T1-weighted structural images. The resulting functional images were band-pass filtered (0.008Hz < f < 0.09Hz).

Regional time-series were extracted using the Harvard-Oxford cortical and subcortical structural atlas (48 cortical regions: 43 bilateral, 5 others; 25 subcortical regions: 16 bilateral, 9 others; totaling 132 = 43 × 2 + 5 + 16 × 2 + 9). This parcellation was chosen based on previous recommendations (McCulloch et al., 2022) to ensure comparability with other analyses (Müller et al., 2017) and evaluate the face validity of the rDCM approach.

#### 4.4.2 Connectivity analyses

##### Functional Connectivity

We computed functional connectivity using Pearson correlations between the BOLD signal time series of each pair of distinct brain regions. The full functional connectivity matrix of all 132 × 132 regions contained 17424 connections. Diagonal and off-diagonal entries in the upper triangle were not considered in subsequent analyses, yielding 8648 unique correlation coefficients.

##### Effective Connectivity

EC was estimated from the raw timeseries of all 132 regions of interest (ROIs) using rDCM (Frässle et al., 2017, 2018, 2021b). rDCM is a reformulation of the original DCM for fMRI framework (Friston et al., 2003). In general, DCM describes the brain as a non-linear dynamic system, which can be summarized by two equations: a neural state equation and a (non-linear) observer equation. In classic DCM, the state equation describes how activity in brain regions evolves over time, as a function of endogenous connectivity between brain regions, changes in connectivity and activity induced by experimental inputs. In this study, we modelled task-free data and did not consider external experimental inputs. The observer equation describes how changes in neuronal states translate into changes in measurements. For fMRI data, this is a non-linear hemodynamic model (Buxton et al., 1998; Friston et al., 2000; Havlicek et al., 2015; Stephan et al., 2007). Under suitable assumptions and choice of prior distributions over the parameters, the likelihood function can be specified and the model inverted to estimate effective connectivity from fMRI data.

In classic DCM, analyses need to be restricted to model networks of only a few regions using experiments, which carefully control experimental perturbations of network activity to ensure model invertibility (Frässle et al., 2017). To meet these practical constraints *a priori* knowledge of the system under analysis is required. In contrast, rDCM enables inference of EC in whole-brain networks. Through a series of simplifications including, crucially, the transformation into the frequency-(rather than time-) domain, the original DCM model can be recast as a linear regression in the frequency-domain (hence regression or rDCM) reducing its concomitant computational burden and overcoming some of the limitations of classic DCM (Friston et al., 2003):

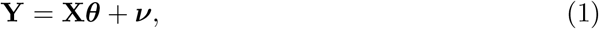

with

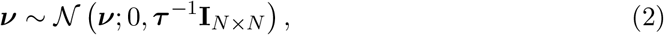

where **Y** is the independent variable (proportional to the Fourier-transformed BOLD signals **ŷ**_i_); **X** represents the design matrix, which consists of the regressors (the Fourier-transformed BOLD signals); ***θ*** is the parameter vector, which consists of the endogenous connectivity strengths *a_i_j* from region *i* to region *j,* and ***ν*** is the noise, with precision 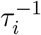 distinct for each region.

Additional simplifications from the original DCM framework include: (i) replacing the nonlinear hemodynamic model with a linear hemodynamic response function, (iii) applying a mean field approximation across regions (i.e., connectivity parameters targeting different regions are assumed to be independent), and (iv) specifying conjugate priors on neuronal (i.e., connectivity and driving input) parameters and noise precision. The face- and construct-validity of rDCM for use with rs-fMRI data was recently demonstrated by (Frässle et al., 2021b). When we refer to EC, we mean the estimated endogenous connectivity strength parameter values *a_ij_*. rDCM parameters were estimated for fully connected whole-brain networks using the rDCM toolbox, which is made available as open-source code as part of the TAPAS (Frässle et al., 2021a) software collection (version 4.4.0; http://www.translationalneuromodeling.org/tapas).

#### 4.4.3 Statistical analysis

We investigated connectivity changes under LSD using three complementary approaches: (1) mass-univariate tests; (2) multivariate tests; and (3) machine learning, which allowed us to assess multi-variate, non-linear changes and quantify how well we were able to distinguish LSD from placebo states at the single-participant level. This last analysis served to gauge the potential of using connectivity features to predict subjective effects or clinical outcomes in future studies.

##### Mass-univariate analysis

We compared changes in connectivity under LSD using mass-univariate (paired-sample) t-tests with Benjamin and Hochberg (Benjamini and Hochberg, 1995) correction to adjust for multiple testing and control the false discovery rate (FDR) of a family of hypothesis test, which was set to *α* < 0.05. Multiple testing correction was implemented using the fdr_bh function ((Groppe, 2015); https://www.mathworks.com/matlabcentral/fileexchange/27418-fdr_bh).

##### Partial least squares correlation analysis

Partial least squares correlation (PLSC) analysis was used to capture connections that maximally represented differences between LSD and placebo. PLS is similar to principal components analysis (PCA) or canonical correlation, with the exception of one important feature: PLS solutions are constrained to the part of the covariance structure that is attributable to the experimental manipulations (McIntosh and Lobaugh, 2004).

Singular value decomposition (SVD) was applied to the mean-centered matrix of FC, EC, and self-connections separately. Mathematically, SVD simply re-expresses this matrix as a set of orthogonal singular vectors or latent variables (LVs), the number of which is equivalent to the total number of conditions. The LVs can be understood analogous to principal components in PCA and account for the covariance of the original mean-centered matrix in decreasing order of magnitude. Statistical significance of the obtained LVs and reliability of region loadings on the LVs were assessed through permutation tests with 2000 permutations bootstrapping with 2000 samples, respectively.

##### Machine learning analysis

To assess how well LSD could be distinguished from placebo at the individual level we trained two random forest classifiers (Breiman, 2001)on either whole-brain FC or EC as features to predict condition. The number of decision trees was kept constant and equivalent to the number of features for each model without further optimization (due to computational constraints and the high-dimensionality of the data). All other parameters were kept to the default values. Both classifiers were implemented using the ‘TreeBagger’ function.

Feature preprocessing consisted of (i) pruning and (ii) covariate correction (Koutsouleris et al., 2012) for baseline heart rate, diastolic and systolic blood pressure, and body temperature, and was embedded in a 5-fold cross-validation. Since data were collected using a within-subject design, we additionally ensured that both conditions of an individual were included either in the training *or* the test set to exclude that generalisibility estimates were influenced by person-specific information and prevent potential information leakage. We report accuracy (ACC), balanced accuracy (BAC), sensitivity (SE), specificity (SP), positive and negative predictive values (PPV and NPV), and area-under-the-curve (AUC) to evaluate classification performances. Classification performances wer tested against chance performance using permutation tests with 1000 label permutations and FC vs EC classifiers were compared against each other using McNemar’s tests.

Lastly, feature importance was computed based on the mean decrease in accuracy on permuted out-of-bag samples for each connection. This measure indicates how much performance decreases when removing a feature from the model and reflects the importance of this feature for the overall classification performance (a larger decrease in accuracy indicates higher importance).

## 5 Acknowledgements

We are grateful for support by the Swiss National Science Foundation (trial A: SNF project grant, 320030_1449493 to MEL, trial B: SNF project grant, 320030_170249 to SB and MEL, Doc.Mobility, P1BSP3_200054 to DJH, Ambizione, PZ00P_167952 to AOD) and the Krembil Foundation (to AOD).

## References

Aghajanian, G. K. and Marek, G. J. (1999). Serotonin, via 5-HT2A receptors, increases EPSCs in layer V pyramidal cells of prefrontal cortex by an asynchronous mode of glutamate release. Brain Research, 825(1-2):161–171.

Ashburner, J. and Friston, K. (2005). Unified segmentation. NeuroImage, 26(3):839–51.

Barnett, L., Muthukumaraswamy, S. D., Carhart-Harris, R. L., and Seth, A. K. (2020). Decreased directed functional connectivity in the psychedelic state. NeuroImage, 209(116462).

Beliveau, V., Ganz, M., Feng, L., Ozenne, B., Højgaard, L., Fisher, P. M., Svarer, C., Greve, D. N., and Knudsen, G. M. (2017). A high-resolution in vivo atlas of the human brain’s serotonin system. Journal of Neuroscience, 37(1):120–128.

Benjamini, Y. and Hochberg, Y. (1995). Controlling the false discovery rate: A practical and powerful approach to multiple testing. Journal of the Royal Statistical Society, Series B (Methodological), 57(1):289–300.

Bershad, A. K., Preller, K. H., Lee, R., Keedy, S., Wren-Jarvis, J., Bremmer, M. P., and de Wit, H. (2020). Preliminary report on the effects of a low dose of LSD on restingstate amygdala functional connectivity. Cognitive neuroscience and neuroimaging, 5(4):461–467.

Birn, R., Molloy, E., Patriat, R., Parker, T., Meier, T., Kirk, G., Nair, V., Meyerand, M., and Prabhakaran, V. (2013). The effect of scan length on the reliability of resting-state fMRI connectivity estimates. NeuroImage, 83:550–8.

Bogenschutz, M. P., Ross, S., Bhatt, S., Baron, T., Forcehimes, A. A., Laska, E., Mennenga, S. E., O’Donnell, K., Owens, L. T., Podrebarac, S., et al. (2022). Percentage of heavy drinking days following psilocybin-assisted psychotherapy vs placebo in the treatment of adult patients with alcohol use disorder: a randomized clinical trial. JAMA Psychiatry, 79(10):953–962.

Breiman, L. (2001). Random forests. Machine Learning, 45:5–32.

Buxton, R., Wong, E., and Frank, L. (1998). Dynamics of blood flow and oxygenation changes during brain activation: the balloon model. Magnetic Resonance in Medicine., 39(6):855–864.

Carhart-Harris, R., Giribaldi, B., Watts, R., Baker-Jones, M., Murphy-Beiner, A., Murphy, R., Martell, J., Blemings, A., Erritzoe, D., and Nutt, D. J. (2021). Trial of psilocybin versus escitalopram for depression. New England Journal of Medicine, 384(15):1402–1411.

Carhart-Harris, R. L. (2018). The entropic brain-revisited. Neuropharmacology, 142:167–178.

Carhart-Harris, R. L., Bolstridge, M., Day, C., Rucker, J., Watts, R., Erritzoe, D., Kaelen, M., Giribaldi, B., Bloomfield, M., Pilling, S., et al. (2018). Psilocybin with psychological support for treatment-resistant depression: six-month follow-up. Psychopharmacology, 235(2):399–408.

Carhart-Harris, R. L., Bolstridge, M., Rucker, J., Day, C. M., Erritzoe, D., Kaelen, M., Bloomfield, M., Rickard, J. A., Forbes, B., Feilding, A., et al. (2016a). Psilocybin with psychological support for treatment-resistant depression: an open-label feasibility study. The Lancet Psychiatry, 3(7):619–627.

Carhart-Harris, R. L. and Friston, K. J. (2019). Rebus and the anarchic brain: toward a unified model of the brain action of psychedelics. Pharmacological Reviews, 71(3):316–344.

Carhart-Harris, R. L., Leech, R., Hellyer, P. J., Shanahan, M., Feilding, A., Tagliazucchi, E., Chialvo, D. R., and Nutt, D. (2014). The entropic brain: a theory of conscious states informed by neuroimaging research with psychedelic drugs. Frontiers in Human Neuroscience, 8.

Carhart-Harris, R. L., Muthukumaraswamy, S., Roseman, L., Kaelen, M., Droog, W., Murphy, K., Tagliazucchi, E., Schenberg, E. E., Nest, T., Orban, C., Leech, R., Williams, L. T., Williams, T. M., Bolstridge, M., Sessa, B., McGonigle, J., Sereno, M. I., Nichols, D., Hellyer, P. J., Hobden, P., and Nutt, D. J. (2016b). Neural correlates of the LSD experience revealed by multimodal neuroimaging. Proceedings of the National Academy of Sciences of the United States of America, 113(17):4853–4858.

Davis, A. K., Barrett, F. S., May, D. G., Cosimano, M. P., Sepeda, N. D., Johnson, M. W., Finan, P. H., and Griffiths, R. R. (2021). Effects of psilocybin-assisted therapy on major depressive disorder: a randomized clinical trial. JAMA Psychiatry, 78(5):481–489.

Dolder, P. C., Liechti, M. E., and Rentsch, K. M. (2018). Development and validation of an lc-ms/ms method to quantify lysergic acid diethylamide (lsd), iso-lsd, 2-oxo-3-hydroxy-lsd, and nor-lsd and identify novel metabolites in plasma samples in a controlled clinical trial. Journal of Clinical Laboratory analysis, 32(2):e22265.

Dolder, P. C., Schmid, Y., Müller, F., Borgwardt, S., and Liechti, M. E. (2016). Lsd acutely impairs fear recognition and enhances emotional empathy and sociality. Neuropsychopharmacology, 41(11):2638–2646.

Felleman, D. J. and Van Essen, D. C. (1991). Distributed hierarchical processing in the primate cerebral cortex. Cerebral Cortex, 1(1):1–47.

Friston, K. (1994). Functional and effective connectivity in neuroimaging: A synthesis. Human Brain Mapping, 2(1-2).

Friston, K., Mechelli, A., Turner, R., and Price, C. (2000). Nonlinear responses in fMRI: the balloon model, volterra kernels and other hemodynamics. NeuroImage, 12(4):466–477.

Friston, K. J., Ashburner, J., Kliebel, S., Nichols, T., and Penny, W. (2007). Statistical Parametric Mapping: The Analysis of Functional Brain Images. Elsevier, 1 edition.

Friston, K. J., Harrison, L., and Penny, W. (2003). Dynamic causal modelling. NeuroImage, 19(4):1273–1302.

Frässle, S., Aponte, E., Bollmann, S., Brodersen, K., Do, C., Harrison, O., Harrison, S., Heinzle, J., Iglesias, S., Kasper, L., Lomakina, E., Mathys, C., Müller-Schrader, M., Pereira, I., Petzschner, F., Raman, S., Schöbi, D., Toussaint, B., Weber, L., Yao, Y., and Stephan, K. (2021a). TAPAS: an open-source software package for translational neuromodeling and computational psychiatry. Frontiers in Psychiatry, 12(857).

Frässle, S., Harrison, S. J., Heinzle, J., Clementz, B. A., Tamminga, C. A., Sweeney, J. A., Gershon, E. S., Keshavan, M. S., Pearlson, G. D., Powers, A., and Stephan, K. E. (2021b). Regression dynamic causal modeling for resting-state fMRI. Human Brain Mapping, 42(7):2159–2180.

Frässle, S., Lomakina, E. I., Kasper, L., M., M. Z., Leff, A., Pruessmann, K. P., Buhmann, J. M., and Stephan, K. E. (2018). A generative model of whole-brain effective connectivity. NeuroImage, 179(505-529).

Frässle, S., Lomakina, E. I., Razi, A., Friston, K. J., Buhmann, J. M., and Stephan, K. E. (2017). Regression DCM for fMRI. NeuroImage, 155:406–421.

Gasser, P., Kirchner, K., and Passie, T. (2015). LSD-assisted psychotherapy for anxiety associated with a life-threatening disease: a qualitative study of acute and sustained subjective effects. Journal of Psychopharmacology, 29(1):57–68.

Girn, M., Roseman, L., Bernhardt, B., Smallwood, J., Carhart-Harris, R., and Spreng, R. (2022). Serotonergic psychedelic drugs LSD and psilocybin reduce the hierarchical differentiation of unimodal and transmodal cortex. NeuroImage, 256(119220).

Griffiths, R. R., Johnson, M. W., Carducci, M. A., Umbricht, A., Richards, W. A., Richards, B. D., Cosimano, M. P., and Klinedinst, M. A. (2016). Psilocybin produces substantial and sustained decreases in depression and anxiety in patients with lifethreatening cancer: A randomized double-blind trial. Journal of Psychopharmacology, 30(12):1181–1197.

Griffiths, R. R., Richards, W. A., McCann, U., and Jesse, R. (2006). Psilocybin can occasion mystical-type experiences having substantial and sustained personal meaning and spiritual significance. Psychopharmacology, 187(3):268–283.

Groppe, D. (2015). fdr_bh. https://www.mathworks.com/matlabcentral/fileexchange/27418-fdr_bh), MATLAB Central File Exchange. Retrieved 08-03-2021.

Havlicek, M., Roebroeck, A., Friston, K., Gardumi, A., Ivanov, D., and Uludag, K. (2015). Physiologically informed dynamic causal modeling of fMRI data. NeuroImage, 122:355–372.

Holze, F., Gasser, P., Müller, F., Dolder, P. C., and Liechti, M. E. (2022). Lysergic acid diethylamide-assisted therapy in patients with anxiety with and without a life-threatening illness a randomized, double-blind, placebo-controlled Phase II study. Biological Psychiatry.

Holze, F., Vizeli, P., Muäller, F., Ley, L., Duerig, R., Varghese, N., Eckert, A., Borgwardt, S., and Liechti, M. (2020). Distinct acute effects of LSD, MDMA and D-amphetamine in healthy subjects. Neuropsychopharmacology, 45(3):462–471.

Jardri, R. and Deneve, S. (2013). Circular inferences in schizophrenia. Brain, 136(11):3227–3241.

Jardri, R., Duverne, S., Litvinova, A. S., and Denève, S. (2017). Experimental evidence for circular inference in schizophrenia. Nature communications, 8(14218):1–13.

Kaelen, M., Roseman, L., Kahan, J., Santos-Ribeiro, A., Orban, C., Lorenz, R., Barrett, F. S., Bolstridge, M., Williams, T., Williams, L., Wall, M. B., Feilding, A., Muthukumaraswamy, S., Nutt, D. J., and Carhart-Harris, R. (2016). LSD modulates music-induced imagery via changes in parahippocampal connectivity. European Neuropsychopharmacology: The Journal of the European College of Neuropsychopharmacology, 26(7):1099–1109.

Kasper, L., Bollmann, S., Diaconescu, A. O., Hutton, C., Heinzle, J., Iglesias, S., Hauser, T. U., Sebold, M., Manjaly, Z.-M., Pruessmann, K. P., et al. (2017). The physio toolbox for modeling physiological noise in fmri data. Journal of Neuroscience Methods, 276:56–72.

Koutsouleris, N., Borgwardt, S., Meisenzahl, E., Bottlender, R., Möller, H., and Riecher-Rässler, A. (2012). Disease prediction in the at-risk mental state for psychosis using neuroanatomical biomarkers: results from the FePsy study. Schizophrenia Bulletin, 38(6):1234–46.

Krebs, T. S. and Johansen, P.-Ø. (2012). Lysergic acid diethylamide (LSD) for alcoholism: meta-analysis of randomized controlled trials. Journal of Psychopharmacology, 26(7):994–1002.

Leptourgos, P., Bouttier, V., Denève, S., and Jardri, R. (2022). From hallucinations to synaesthesia: a circular inference account of unimodal and multimodal erroneous percepts in clinical and drug-induced psychosis. Neuroscience & Biobehavioral Reviews, page 104593.

Lewis, C. R., Preller, K. H., Kraehenmann, R., Michels, L., Staempfli, P., and Vollenweider, F. X. (2017). Two dose investigation of the 5-ht-agonist psilocybin on relative and global cerebral blood flow. NeuroImage, 159:70–78.

Luppi, A. I., Carhart-Harris, R. L., Roseman, L., Pappas, I., Menon, D. K., and Stamatakis, E. A. (2021). LSD alters dynamic integration and segregation in the human brain. NeuroImage, 227(117653).

Markov, N. T., Ercsey-Ravasz, M. M., Ribeiro Gomes, A., Lamy, C., Magrou, L., Vezoli, J., Misery, P., Falchier, A., Quilodran, R., Gariel, M.-A., et al. (2014). A weighted and directed interareal connectivity matrix for macaque cerebral cortex. Cerebral Cortex, 24(1):17–36.

Mason, N., Kuypers, K., Müller, F., Reckweg, J., Tse, D., Toennes, S., Hutten, N., Jansen, J., Stiers, P., Feilding, A., et al. (2020). Me, myself, bye: regional alterations in glutamate and the experience of ego dissolution with psilocybin. Neuropsychopharmacology, 45(12):2003–2011.

McCulloch, D. E.-W., Knudsen, G. M., Barrett, F. S., Doss, M. K., Carhart-Harris, R. L., Rosas, F. E., Deco, G., Kringelbach, M. L., Preller, K. H., Ramaekers, J. G., Mason, N. L., Müller, F., and Fisher, P. M. (2022). Psychedelic resting-state neuroimaging: A review and perspective on balancing replication and novel analyses. Neuroscience & Biobehavioral Reviews, 138:104689.

McIntosh, A. and Lobaugh, N. (2004). Partial least squares analysis of neuroimaging data: applications and advances. NeuroImage, 23:Suppl 1:S250–63.

Müller, F., Dolder, P. C., Schmidt, A., Liechti, M. E., and Borgwardt, S. (2018). Altered network hub connectivity after acute LSD administration. NeuroImage Clinical, 18:694–701.

Müller, F., Lenz, C., Dolder, P., Lang, U., Schmidt, A., Liechti, M., and Borgwardt, S. (2017). Increased thalamic resting-state connectivity as a core driver of LSD-induced hallucinations. Acta Psychiatrica Scandinavica, 136(6):648–657.

Nichols, D. E. (2016). Psychedelics. Pharmacological Reviews, 68(2):264–355.

Nierhaus, T. and Schön, T., Becker, R., Ritter, P., and Villringer, A. (2009). Background and evoked activity and their interaction in the human brain. Magnetic Resonance Imaging, 27:1140–1150.

Nutt, D. and Carhart-Harris, R. (2021). The current status of psychedelics in psychiatry. JAMA Psychiatry, 78(2):121–122.

Power, J., Barnes, K., Snyder, A., Schlaggar, B., and Petersen, S. (2012). Spurious but systematic correlations in functional connectivity mri networks arise from subject motion. NeuroImage, 59(3):2142–54.

Preller, K., Burt, J., Ji, J., Schleifer, C., Adkinson, B., Stämpfli, P., Seifritz, E., Repovs, G., Krystal, J., Murray, J., Vollenweider, F., and Anticevic, A. (2018). Changes in global and thalamic brain connectivity in LSD-induced altered states of consciousness are attributable to the 5-HT2A receptor. eLife, 7(e35082).

Preller, K., Razi, A., Zeidman, P., Stämpfli, P., Seifritz, E., Friston, K., and Vollenweider, F. (2019). Effective connectivity changes in LSD-induced altered states of consciousness in humans. Proceedings of the National Academy of Sciences of the United States of America, 116(7):2743–2748.

Roseman, L., Nutt, D. J., and Carhart-Harris, R. L. (2018). Quality of acute psychedelic experience predicts therapeutic efficacy of psilocybin for treatment-resistant depression. Frontiers in Pharmacology, 8:974.

Roseman, L., Sereno, M. I., Leech, R., Kaelen, M., Orban, C., McGonigle, J., Feilding, A., Nutt, D. J., and Carhart-Harris, R. L. (2016). LSD alters eyes-closed functional connectivity within the early visual cortex in a retinotopic fashion. Human Brain Mapping, 37(8):3031–3040.

Ross, S., Bossis, A., Guss, J., Agin-Liebes, G., Malone, T., Cohen, B., Mennenga, S. E., Belser, A., Kalliontzi, K., Babb, J., et al. (2016). Rapid and sustained symptom reduction following psilocybin treatment for anxiety and depression in patients with life-threatening cancer: a randomized controlled trial. Journal of Psychopharmacology, 30(12):1165–1180.

Rucker, J. J., Iliff, J., and Nutt, D. J. (2018). Psychiatry & the psychedelic drugs. past, present & future. Neuropharmacology, 142:200–218.

Schmid, Y., Enzler, F., Gasser, P., Grouzmann, E., Preller, K. H., Vollenweider, F. X., Brenneisen, R., Muller, F., Borgwardt, S., and Liechti, M. E. (2015). Acute effects of lysergic acid diethylamide in healthy subjects. Biological Psychiatry, 78(8):544–553.

Scruggs, J. L., Patel, S., Bubser, M., and Deutch, A. Y. (2000). Doi-induced activation of the cortex: dependence on 5-ht2a heteroceptors on thalamocortical glutamatergic neurons. Journal of Neuroscience, 20(23):8846–8852.

Scruggs, J. L., Schmidt, D., and Deutch, A. Y. (2003). The hallucinogen 1-[2, 5-dimethoxy-4-iodophenyl]-2-aminopropane (doi) increases cortical extracellular glutamate levels in rats. Neuroscience letters, 346(3):137–140.

Stephan, K., Weiskopf, N., Drysdale, P., Robinson, P., and Friston, K. (2007). Comparing hemodynamic models with DCM. NeuroImage, 38:387–401.

Tagliazucchi, E., Roseman, L., Kaelen, M., Orban, C., Muthukumaraswamy, S. D., Murphy, K., Laufs, H., Leech, R., McGonigle, J., Crossley, N., Bullmore, E., Williams, T., Bolstridge, M., Feilding, A., Nutt, D. J., and Carhart-Harris, R. (2016). Increased global functional connectivity correlates with LSD-induced ego dissolution. Current Biology, 26(8):1043–1050.

Timmermann, C., Spriggs, M. J., Kaelen, M., Leech, R., Nutt, D. J., Moran, R. J., Carhart-Harris, R. L., and Muthukumaraswamy, S. D. (2018). LSD modulates effective connectivity and neural adaptation mechanisms in an auditory oddball paradigm. Neuropharmacology, 142:251–262.

Vollenweider, F. X. and Geyer, M. A. (2001). A systems model of altered consciousness: integrating natural and drug-induced psychoses. Brain Research Bulletin, 56(5):495–507.

Whitfield-Gabrieli, S. and Nieto-Castanon, A. (2012). Conn: a functional connectivity toolbox for correlated and anticorrelated brain networks. Brain Connectivity, 2(3):125–141.

